# Significance of Oct-4 transcription factor as a pivotal therapeutic target for CD44^+^/24^-^ mammary tumour initiating cells; aiming at the root of the recurrence

**DOI:** 10.1101/2021.05.21.445099

**Authors:** Utsav Sen, Shanooja Shanavas, Apoorva Gowda, Muhammad Nihad, Hari Krishnareddy Rachamalla, Rajkumar Banerjee, Sudheer Shenoy P, Bipasha Bose

## Abstract

Breast cancer (BC) remains one of the deadliest and frequently diagnosed metastatic cancers worldwide. Cancer stem cells (CSCs) are the cell population within the tumour niche, having an epithelial to mesenchymal (EMT) transition phenotype, high self-renewal, vigorous metastatic capacity, drug resistance, and tumour relapse. Identification of targets for induction of apoptosis is essential to provide novel therapeutic approaches in BC. Our earlier studies showed that Vitamin C induces apoptotic cell death by losing redox balance in TNBC CSCs. In this study, we have attempted to identify previously unrecognized CSC survival factors that can be used as druggable targets for bCSCs apoptosis regulators isolated from the TNBC line, MDA MB 468. After a thorough literature review, Oct-4 was identified as the most promising marker for its unique abundance in cancer and absence in normal cells and the contribution of Oct-4 to the sustenance of cancer cells. We then validated a very high expression of Oct-4 in the MDA MB 468 bCSCs population using flow-cytometry. The loss of Oct-4 was carried out using siRNA-mediated knockdown in the bCSCs, followed by assessing for cellular apoptosis. Our results indicated that Oct-4 knockdown induced cell death, changes in cellular morphology, inhibited mammosphere formation, and positive for Annexin-V expression, thereby indicating the role of Oct-4 in bCSC survival. Moreover, our findings also suggest the direct interaction between Oct-4 and Vitamin C using *in silico* docking. This data, hence, contributes towards novel information about Oct-4 highlighting this molecule as a novel survival factor in bCSCs.

## 1. Introduction

Triple-negative breast cancer (TNBC) is a notable clinical problem to which 30% of all patients often turn up with a recurrence of the disease within 2 to 5 years after completion of treatment (Yin et al., 2020). Despite extensive progress in BC’s diagnosis and treatment, several clinical and scientific problems remain unsolved (Harbeck et al., 2019). Procedures for advanced stages of this disease are still relatively limited and inefficient (Mattina et al., 2016). The limitation of these therapies is due to not yet efficiently targeting two essential events that are happening to the breast cancer cells, i.e., epithelial to mesenchymal transition (EMT) and cancer stem cells (CSCs) turnover (Saadin & White 2013).

Based on pathology and gene expression profiling, triple-negative breast cancer cells (ER^−^, PR^−^, HER2^−^) (TNBCs) are heterogeneous and enriched with CD44^+^/24^-^ tumour-initiating cells or cancer stem cells (TICs/CSCs) (Collina et al., 2015, Das et al., 2017). These pathobiological modifications make TNBC cells aggressive, metastatic, and less sensitive to standard chemotherapy and tumour relapse. These cancer stem cells are interlinked with each other and play crucial roles in BC progression and relapse (Wang et al., 2014; Fabregat, Malfettone & Soukupova et al., 2016; Radosa et al., 2016). For metastasis, cancer cells modify their cytoskeleton structure, thus initiating invasion and migration of the CSCs (Fares et al., 2020). The initiation of the migration invasion and metastasis is associated with EMT (Kristensen et al., 2011; Li et al., 2017). Where the cells lose the epithelial markers like E-cadherin, EpCAM, and gain mesenchymal markers like N-cadherin, Vimentin, Snail, and Twist (Loh et al., 2019).

Homeodomain transcription factor of the Pit-Oct-Unc (POU) family, named Octamer-binding transcription factor 4 (Oct-4) is considered to play an essential role in the self-renewal epithelial-mesenchymal transition (EMT) (Zeineddine et al., 2014) and drug resistance development of CSCs and breast cancer metastasis (Wang et al., 2014). Oct-4 is well established as one of the most important transcription factors that control the self-renewal of pluripotent stem cells and cancer cells (Kim, & Nam 2011). Most interestingly, Oct-4 expression is predominantly observed in embryonic stem cells (ESCs) or some cancer cells (Wang & Herlyn 2015). Hence, this uniqueness of abundance and the crucial developmental and sustenance role could make Oct-4 one of the critical factors worth targeting in stem cell-specific cancer therapy (Wu & Schöler 2014). In our previous study, we have used Vitamin C, and demonstrated the apoptotic effect of Vitamin C on bCSCs (Liu, Yu & Liu 2013).

In this study, we have first isolated the breast CSCs from the TNBC cells MDA MB 468, using flow-sorting based on surface marker CD44^+^/CD24^-^. Then the abundance of the Oct-4 was confirmed by using flow cytometric analysis in bCSCs. Loss of function of Oct-4 on bCSCs survival was established upon Oct-4 knockdown using siRNA. In summary, we have deciphered the importance of the transcription factor Oct-4 in bCSC survival and sustenance. Also, we sought to determinize the efficiency of Oct-4 as a potential target for the bCSCs toward cancer/cancer stem cell therapeutics.

### 2.0 Material and Methods

### 2.1 Cell culture

The TNBC line MDA MB 468 was procured from the National Centre for Cell Science, Pune, India and the NTERA-2 CLD1 cells were purchased from ATCC, USA. The cancer cells were cultured in media containing DMEM supplemented with 10% fetal bovine serum (Cat. No. RM9955) (all from Hi-Media Laboratories, India), 1% Penicillin and Streptomycin, GlutaMAX, CSCs and NTERA -2 cells were supplemented with non-essential amino acids, sodium pyruvate, and 0.1% of 2-mercaptoethanol (all from Gibco Thermo Fisher Scientific, USA). The cells were grown in incubator with 5% CO_2_ with humidified atmosphere. The cells were trypsinized upon reaching 75–80% confluency using 0.25% trypsin-EDTA (Thermo Fisher Scientific, USA).

### 2.2 Immunophenotyping by flow cytometry

For analytical flow cytometry (immunophenotyping), the bCSCs were fixed with 4% PFA and permeabilized with 0.3% Triton X-100. Cells were then incubated with the fluorescently tagged antibody (CD44 and CD24) for 1 hour. Then the cells were given PBS wash and acquired on guava EasyCyte flow cytometer (10,000 events). Obtained data were analyzed using DE Novo FCS Express 5 software, USA (antibody details are available in Supplementary Table-1).

### 2.3 Migration assay

Sorted and MDA MB 468 CSC, and the WT cells were seeded in a 6-well plate. After reaching 80% confluency, the cell monolayer was scraped/scratch using a 100µl pipette tip. The cells were washed with PBS to remove all single cells. Phase-contrast images (10X magnifications) were captured after 24h of the scratch for two cell types. Percentage cell migration was calculated by measuring the distance between the scratches comparing the final gap width to the initial gap width at Time T0 (zero) using Image-J software. Covered area was calculated in 5 random fields and represented as percentage of area covered. Experiments were performed in triplicate.

### 2.4 Mammosphere assay

Sorted and MDA MB 468 CSC and the WT cells were harvested by trypsinization. Cells were counted and around 800 cells per 20μl drops were kept as hanging drops on the lid of 100mm dish at 37°C under humidified conditions. 3D mammosphere of 200-250μm (average diameter) size were formed at the end of 48h for all the four cell types. At the end of four days (96h), mammospheres were collected and observed under the microscope (Primovert inverted microscope from CARL ZEISS) and images were captured. Relative area, roundness score and solidity of 3D mammosphere were calculated using Image-J software.

### 2.5 Immunocytochemistry assay

The cells were cultured on 8 well chamber slides in 70% confluency followed by fixing using 4% paraformaldehyde (PFA) and permeabilization with 0.3% Triton X-100 (Sigma) in PBS. The non-specific binding sites were blocked with 5% fetal bovine serum in PBS (Himedia Laboratories, India) for 1 hour. Cells were then incubated with primary antibodies overnight at 4°C, followed by the appropriate secondary antibody for 1 hour, counterstained with DAPI for 10 minutes. Finally, the slides were mounted with Prolonged Glass Antifade (Thermo Fisher, USA). The slides were observed, and images were captured under a ZOE Fluorescent cell imager (Bio-Rad, USA). The details of the antibodies are provided in (Supplementary table-1).

### 2.6 Gene expression study by qRT-PCR

One million (1×10^6^) cells were used to isolate RNA using TRIzol reagent (15596026, Thermo Scientific, USA) as per the manufacturer’s instructions. Quantification of RNA was done by using a Colibri Micro volume Spectrometer (Titertek-Berthold, Germany). 1μg of RNA was reverse transcribed to cDNA using iSCRIPT™ cDNA synthesis kit (Bio-Rad Catalog Number-1708891), and qRT-PCR reactions were performed using SSO-Fast™ Eva Green Supermix (Bio-Rad, USA) as per the manufacturer’s instructions. The individual mRNA expressions of the tested genes of all the cell types (Ct values) were first normalized with their respective GAPDH values for getting the δCt values. Finally, the respective mRNA expressions were represented as fold change. The primer sequences were designed by the authors (Sigma, United States). The details of primers are provided in (Supplementary table-2)

### 2.7 siRNA transfection

For transfection experiments, 0.5×10^6^ cells were plated in a 6-well plate and allowed to reach 75% confluency. Cells were transfected with 10nM of anti-Oct-4 siRNA and scrambled/control (10 nM) siRNA. D1X cationic lipid-based formulation (developed in CSIR-IICT, Hyderabad) was used as a potent siRNA transfection reagent instead of the commercially available transfection reagents such as lipofectamine or fugene, in the presence of serum containing media as described previously (Rachamalla et al., 2019). After 16h incubation at 37ºC 5% CO_2_, the media was changed, and transfection was confirmed by fluorescence microscopy. Other experiments for validation were done simultaneously. We had initially transfected the Cy5 labelled scrambled siRNA using the lipid-based gene delivery system (D1X) to determine the transfection efficiency which was 100% as assessed using the confocal microscopy.

### 2.8 Annexin-V-FITC assay

Cancer stem cells (0.5 million) control and transfected with siRNA were washed with ice cold PBS and resuspended in annexin-V binding buffer. Cells were incubated with annexin-V conjugated to FITC antibody (Thermo Fisher Scientific, USA) and propidium iodide (PI) for 15 min in dark. Further, the cells were washed and resuspended in annexin-V binding buffer. Cells (10,000 events) were acquired using Guava EasyCyte Flow Cytometer (Millipore Sigma, USA) data analysis was carried out by using DE Novo FCS Express 5 software, USA.

### 2.9 Molecular modelling to study Oct-4 and Vitamin C interactions: Development of protein structure

Protein target Oct-4 structure was downloaded from database Protein Data Bank (PDB) (OCT-4. PDB). Vitamin C structure was downloaded from Pub Chem. (Pub Chem ID54670067) in .sdf format. Conversion of Ligand i.e., Vitamin C from .sdf format to .pdb format was done by using Open Babel (version 2.4.1). Hydrogen atoms and Kollman charges (7.0) were added to the protein by AutoDock MGL-Tool version 1.5.6 and saved as PDBQT file. Hydrogen atoms, Gasteiger charges were added to ligand and were saved in PDBQT format.

#### Protein ligand Interaction

Vitamin C was docked with Oct-4 using Auto Dock 4.2.6. Grid box was obtained by taking inside the entire protein structure (blind docking method). The grid box size maintained in configuration was 36Å × 30Å × 48Å and centre was 57.908, 0.508, and 3.182. All the information like exhaustiveness, coordinates size of grid box was saved in .txt file and assigning the docking report in an out. pdbqt file. Ligand was aligned with protein molecule and analysed for different binding interactions and interacting residues using PyMol. The main purpose of this experiment is to study the mode of interactions and binding site of Vitamin C with Oct-4.

### 2.10 Statistical analysis

All the experiments were carried as biological and technical triplicates and results have been represented as mean ± SD. The differences between more than two groups were analysed by one way analysis of variance (ANOVA) followed by Bonferroni post-hoc test and between two groups student t-test was used. The P values of P≤0.05 (*) and P≤0.01 (**) P≤0.001 (***) were considered as significant. Error bar represents the ± standard error of mean.

## 3. Results

### 3.1 CD44^+^/24^-^ bCSCs expressed EMT genes, exhibited mammosphere forming abilitie

CD44 is considered a potential CSC marker in most cancers, and CD24 is another vital marker whose prognostic value and significance are investigated in combination with CD44 in various cancers, including breast cancer. By using fluorescent activated cells sorting CD44^+^/24^-^ CSCs were isolated from the heterogeneous TNBC cell line MDA MB 468. Almost, 61.70% of the pure CSC population were isolated and utilized in all the further experiments (Fig. 1A). Gene expression of the CD44^+^/24^-^ CSCs was performed by qRT-PCR, the results indicated significant expression (2.5-fold) of stem cell marker CD44 (up-regulation) and CD24 (down-regulation). Less expression of EMT marker such as EpCAM was observed in MDA MB 468 CSCs. The qRT-PCR analysis results also showed up-regulation of metastatic MMP-2, angiogenesis marker SP-1, proliferative marker AKT, perhaps indicating EMT in concordance with increased stemness metastatic nature of CSCs as compared to WT or heterogeneous population (Fig. 1F).

**Figure-1:**
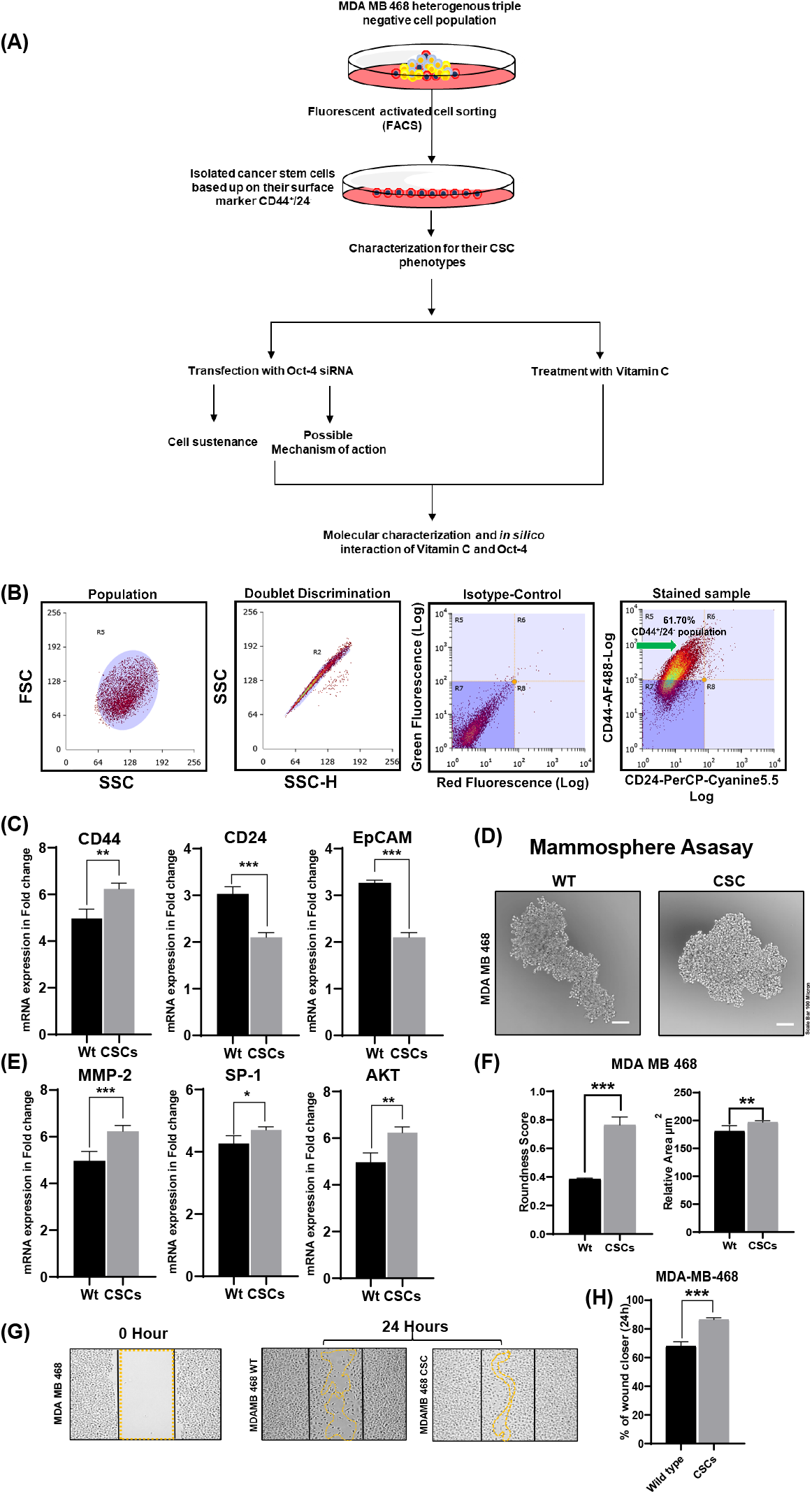
CD44^+^/24^-^ bCSCs from MDA MB 468 TNBC line exhibited CSC phenotype. A) Work flow for the entire work B) FACS sorting of CSCs as CD44^+^ and CD24^-^cells from MDA MB 468. C) and E) Comparative gene expression analysis of both CSCs and WT population from MDA MB468 cells. D) Formation of mammosphere by both Wt. and CSC population F) Quantitative readout of Mammosphere formation by the TNBCs. G) and H) Scratch assay / migration assay results and graphical representation of the data. Scale bar = 100μm. The differences between the two groups were analysed by student t-test. The P values of >0.12 (ns), 0.033(*), 0.002(**), <0.0002(***) were considered as significant. Error bar represents the ± standard error of mean.

In order to test the intrinsic mammosphere forming potential of CD44^+^/24^-^ bCSCs, the mammosphere assay was performed. The mammosphere formed by bCSC were bigger in size and numbers compared to the WT populations (Fig. 1B). The 2D images of the mammospheres upon analysis, exhibited relatively large area (0.570 mm^2^) from CSC as compared that of the mammospheres from WT cells (0.481 mm^2^) (Fig.1C). Furthermore, higher percentage roundness score from CSCs (80%) mammosphere was observed when compared to mammospheres derived from the WT cells (40%) (Fig.1C). These results validated the CSC subpopulation of MDA MB 468, for their self-renewal capacity. In other words, the sorted CSC subpopulation demonstrated more stemness, as compared to, the WT/ parent population.

### 3.2 CD44^+^/24^-^ bCSC population exhibited migratory properties and expression of pluripotent stem cell markers

To validate an in vitro relevance of the migration or metastasis, wound healing assay was performed on CSC populations MDA MB 468 cell line. After 24 hours of the scratch, the distance covered by the cells was highest in the CD44^+^/24^-^ CSCs and less in the WT population (Fig. 1D) indicating a faster migration potential of bCSCs. The percentage of CSC invading the scratch space was more than 80% in MDA MB 468 when compared to WT control (60%) after 24h (Fig.1E).

The stem cell properties of CSCs were also reflected in immunocytochemistry (ICC) analysis. ICC showed that the pluripotent stem cell markers such as Oct-4, Sox-2, Nanog, expression was significantly higher in the CD44^+^/24^-^ bCSCs than the WT/unsorted cells (Fig. 2A and B). The sorted cell types also expressed vimentin suggesting the fact that CSC-like property co-exists with EMT phenotype in bCSC (Wang et al., 2014) (Fig. 2A and B). We have confirmed the pluripotency markers with the positive control NTERA-2 CLD1 cells (Fig. 2C), and compared the expression with the CSCs (Fig. 2D). These results confirmed that the CD44^+^/24^-^ population exhibits CSC characteristics in conjunction with EMT phenotype.

**Figure-2:**
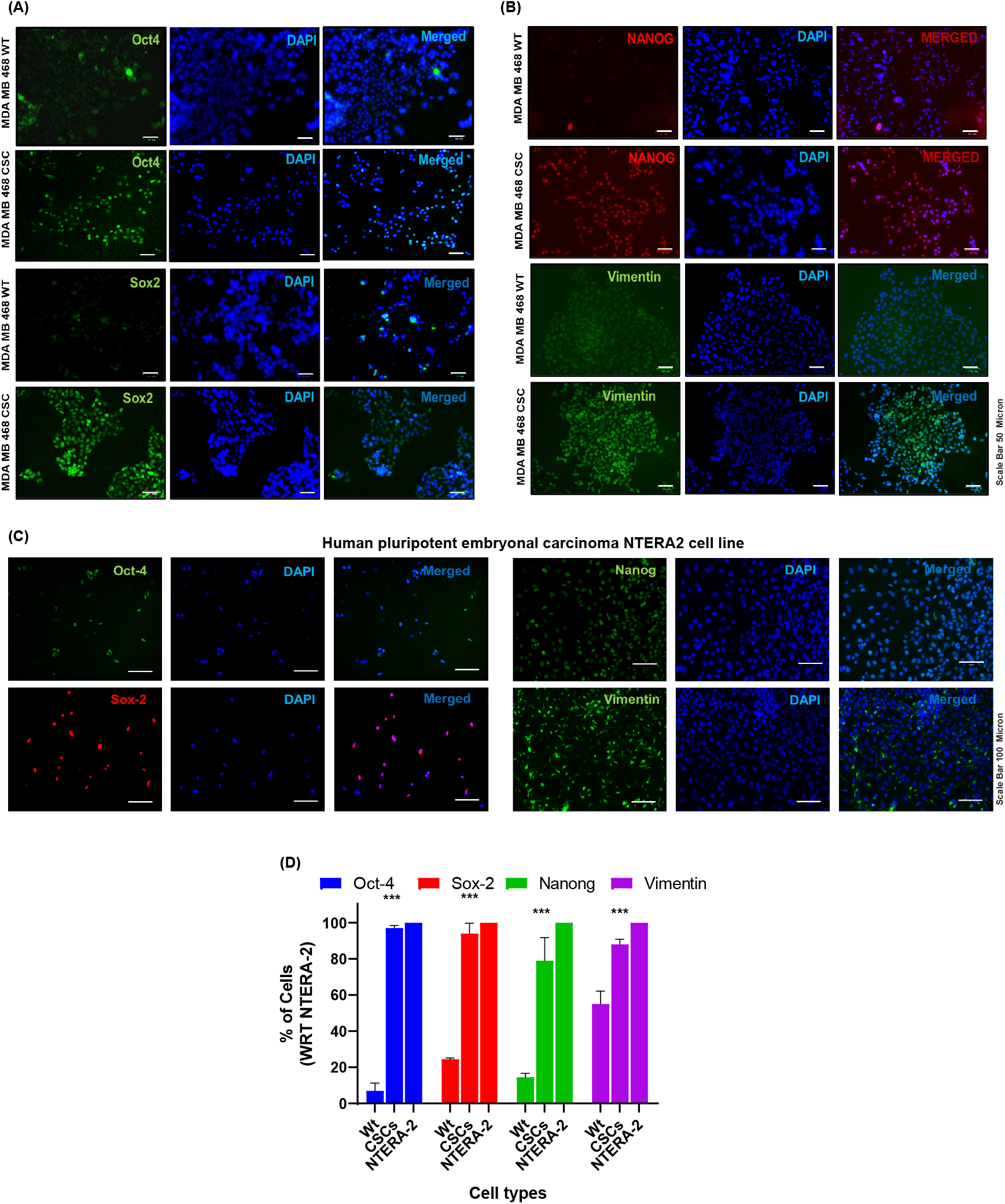
CD44^+^/24^-^ bCSCs from MDA MB 468 TNBC line expressed pluripotency related protein. A) and B) Immunocytochemistry of WT and CSC population from MDAMB 468 line. C) Pluripotency marker expression in NTERA-2 CLD1 pluripotent embryonal carcinoma cell line. D) is the graphical representation of the expression of the percentage of the cells expressing pluripotency marker compared to NTERA-2. Scale bar = 50/100μm. The differences between more than two groups were analysed by one way ANOVA followed by Bonferroni’s post-hoc test. The P values of >0.12 (ns), 0.033(*), 0.002(**), <0.0002(***) were considered as significant. Error bar represents the ± standard error of mean.

### 3.3 Oct-4 plays crucial role in CSCs sustenance and mammosphere formation

Oct-4 is an anti-apoptotic factor and presents itself abundantly in the CSCs to save the cells from programmed cell death (Wang et al., 2013). To assess the importance of the Oct-4 in bCSCs, the Oct-4 was knocked-down using Oct-4 siRNA. When the bCSCs were transfected with the Oct-4 siRNA, as shown in figure-3, the expression level decreased after 16h of transfection (Fig. 3C). Further, the expression level was verified using analytical flow cytometry. In the scrambled sample, 65.23% (Fig.3C) of the cells expressed Oct-4 protein when compared with cells transfected with the Oct-4 siRNA which exhibited only 6-13% (Fig. 3C) of Oct-4 expression. This assay validated the transfection efficiency and downregulation of the Oct-4 expression in bCSCs. Moreover, morphological changes were observed in bCSCs of MDA MB 468 cells after 16 hours post-transfection (Fig.3B). No morphological changes were observed in the cells transfected with Cy5 labelled scrambled siRNA. As Oct-4 has been reported as an anti-apoptotic agent, we hypothesized that siRNA mediated Oct-4 knockdown might induce CSCs apoptosis that was further established by annexin-V expression in the Oct4 knock-down bCSCs (Fig. 3G and H). In summary, knock-down of Oct-4 caused morphological changes along with apoptotic cell-death in bCSCs.

**Figure 3:**
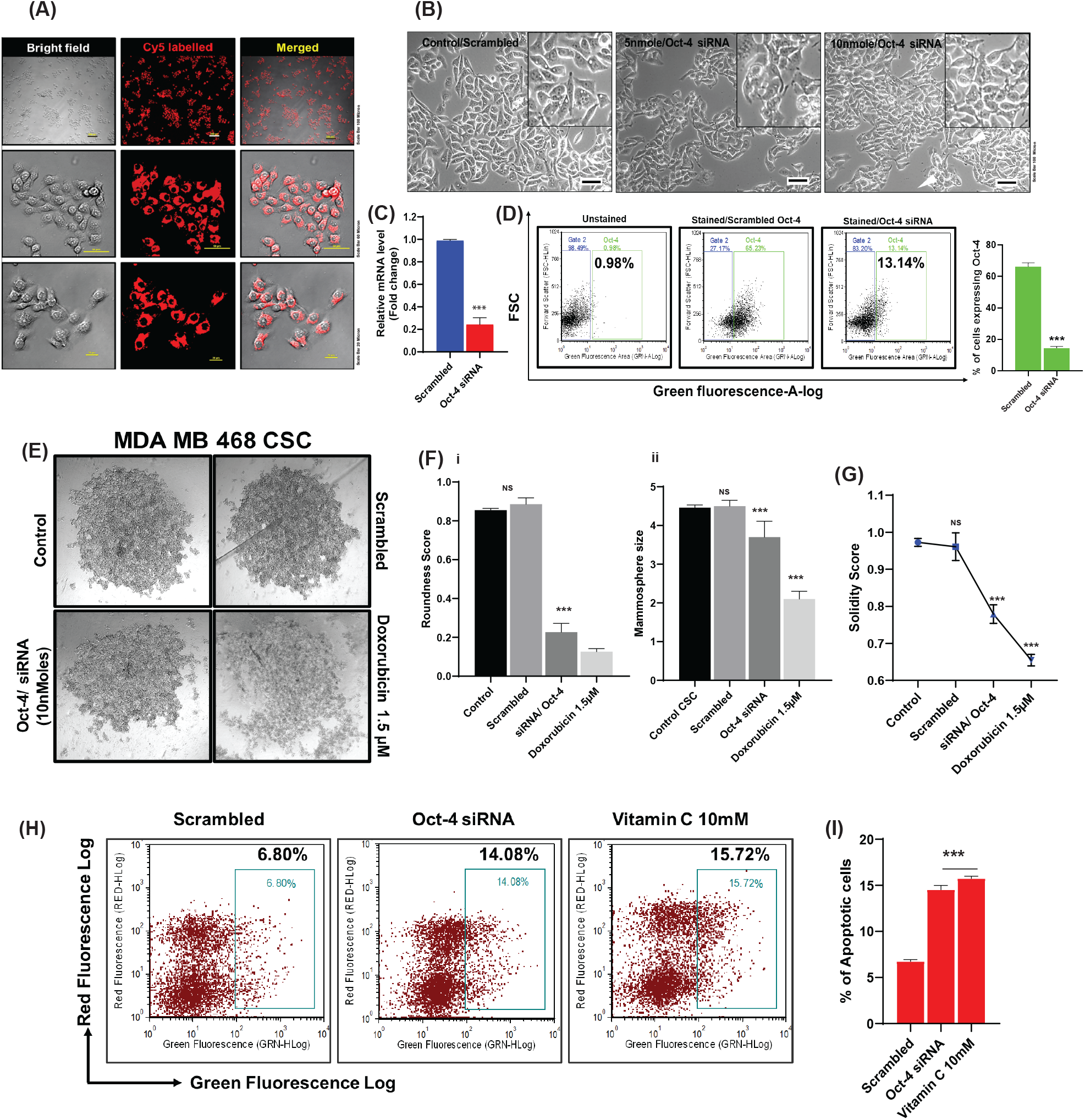
Inhibition of Oct-4 expression prevaricates bCSC survival. A) Confocal images of the MDA MB 468 CSCs after transfected with the Cy5 labelled scrambled siRNA. The Images were captured 16h post-transfection. B) Morphological changes in bCSCs after Oct-4 siRNA transfection C) mRNA expression level of Oct-4 after transfection in mDA-MB-468 sample, compared with the scrambled and siRNA transfected samples D) Analytical flowcytometry result showing the Oct-4 expression before and after transfection. E) Are the representative images obtained from the mammosphere assay after Oct-4 siRNA transfection and scrambled siRNA (Doxorubicin was used as positive control). F) and G) are the direct read out and representation of mammosphere assay. H) Flow cytometry analysis of the Annexin V expression before and after Oct-4 siRNA expression in bCSCs of MDA MB 468 cells, also after treatment with the Vitamin C. I) graphical representation of the percentage of apoptotic cells. Scale bar = 10, 20 and 100μm. The differences between the two groups were analysed by student t-test and more than two groups were analysed by one way ANOVA, followed by Bonferroni’s post-hoc test. The P values of >0.12 (ns), 0.033(*), 0.002(**), <0.0002(***) were considered as significant. Error bar represents the ± standard error of mean.

We next sought to explore the role of Oct-4 in mammosphere formation. As shown in figure 3D, the appearance of the tumour/mammosphere was disrupted in the transfected samples compared to the scrambled sample (Fig. 3D). Both the sphere roundness and the compactness were significantly reduced in the mammospheres formed from si-RNA-Oct4 transfected bCSCs, as compared to, the scrambled transfected bCSCs (Fig. 3D and 3F). Here, 1.5 µM Doxorubicin was used to obtain a positive inhibition of the mammosphere formation. Taken together, the knock-down of Oct-4 caused inhibition of mammosphere formation in bCSCs (Fig. 3D).

### 3.4 Vitamin C directly interacts with bCSCs by down regulation of the pluripotency factor Oct-4

As shown in figure 4A, the bCSCs/CD44^+^24^-^ population when subjected to treated with Vitamin C (10 mM) for 2h, exhibited down-regulation of Oct-4 (Fig. 4A). In the untreated control, the Oct-4 expression was 91.99% in CSCs (Fig. 4A). Interestingly, after Vitamin C treatment, the expression reduced drastically to 3.65% in bCSCs (Figure 4A). As we have mentioned earlier (Sen, Shenoy & Bose et al., 2017, Sen et al., 2020), Vitamin C is responsible for cellular damage, apoptosis, and inhibition of cell proliferation Here we have, observed the down-regulation of the pluripotency-regulating octamer-binding transcription factor (Oct-4) expression upon treatment with Vitamin C for 2h (Fig.4B). Based on the above-mentioned findings, we hypothesized a co-relation/possible interaction between Oct-4 and Vitamin C worth validation.

**Figure 4:**
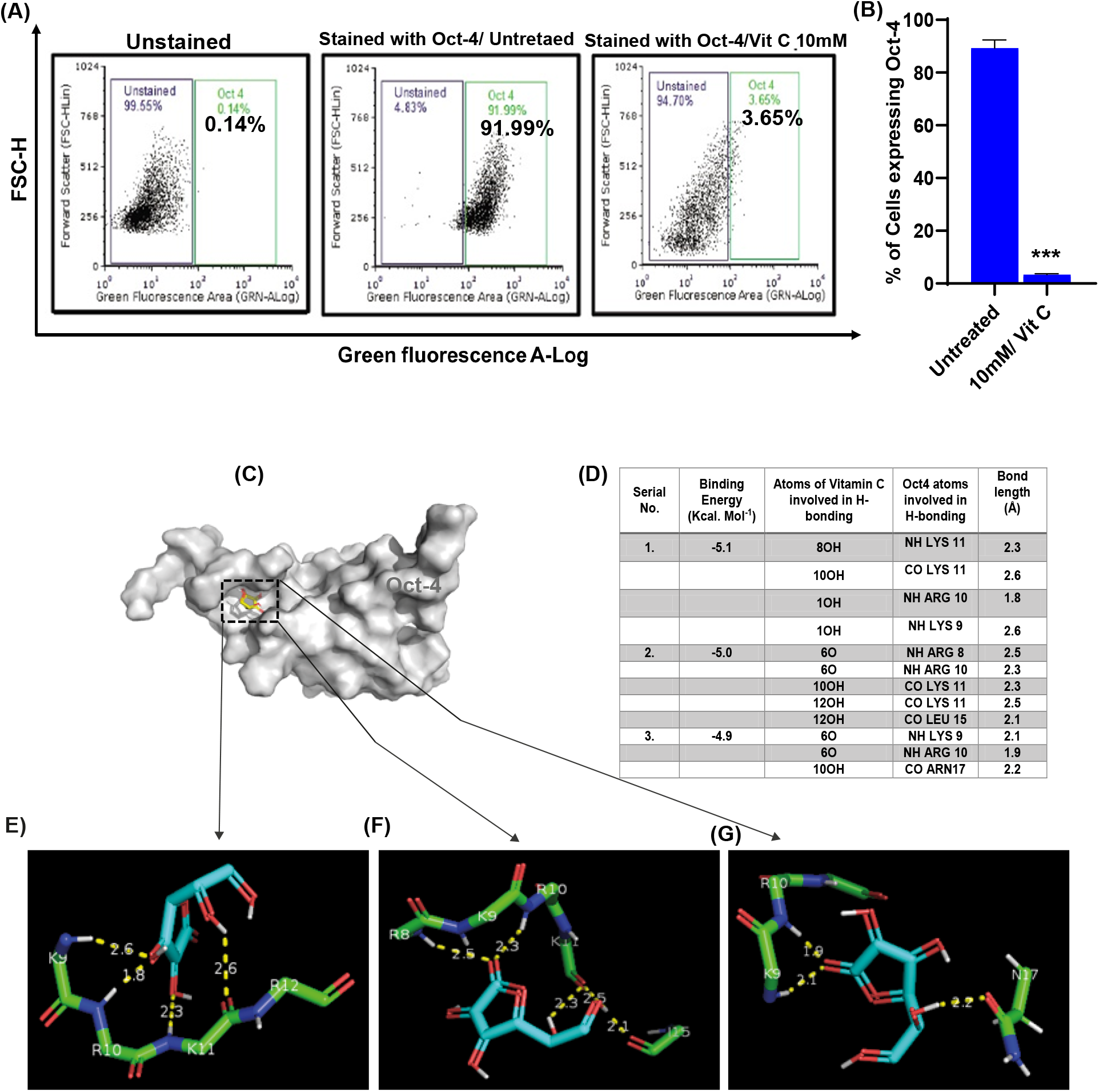
Vitamin C could be a potential target for the Oct-4. Flow cytometry data representation of the Oct-4 expression before and after Vitamin C treatment (10mM). B) graphical representation of the flow data. C)-G) Possible interaction of Vitamin C and Oct-4 protein by *in sillico* docking technique.

**Figure 5:**
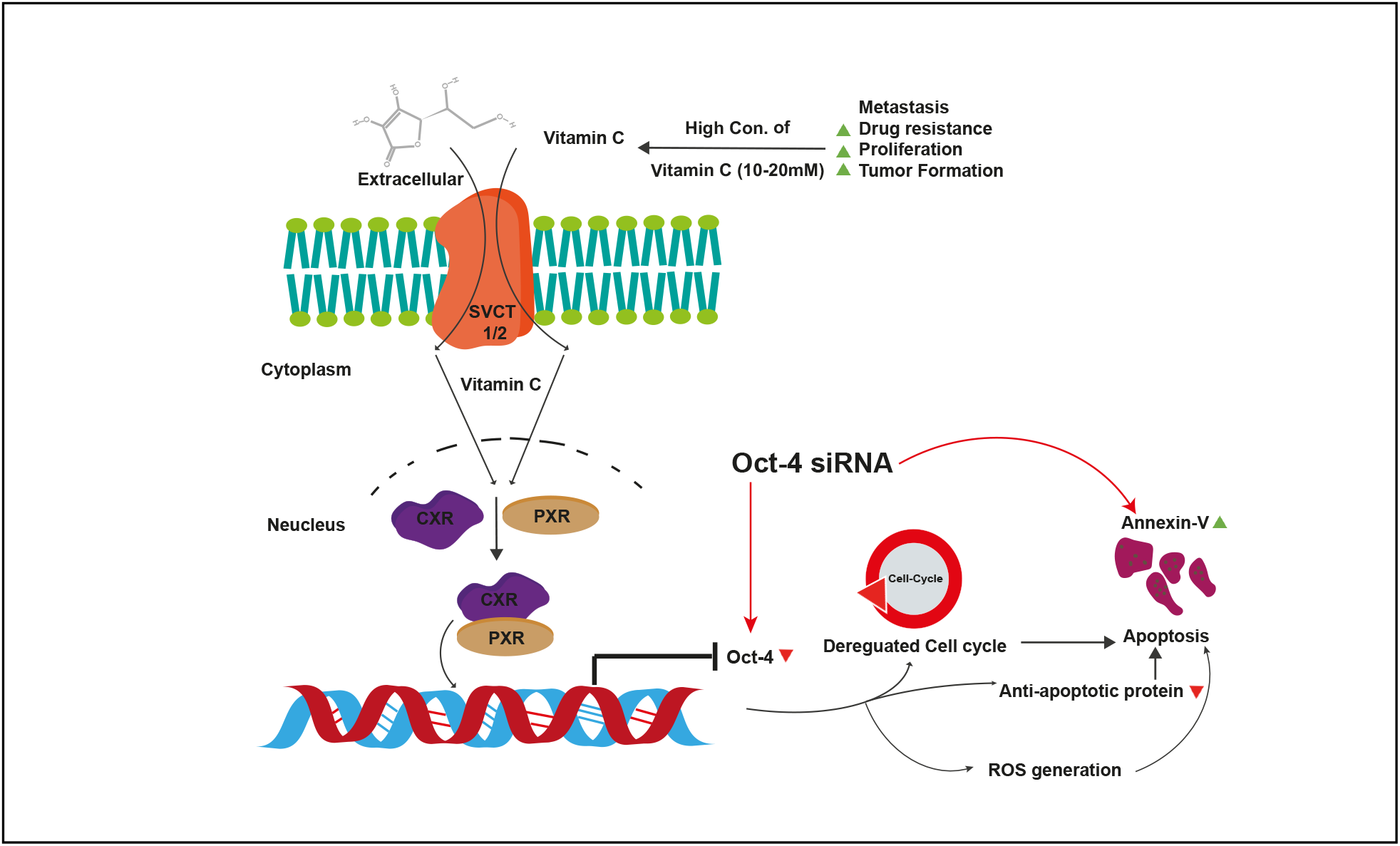
Schematic representation of the work and the possible mechanism of action after Oct-4 knockdown in bCSCs. Oct-4 is very crucial for cell proliferation and sustenance and even for the aggressiveness of cancer. And this unique pluripotent transcription factor is only expressed in ESCs and CSCs. When this transcription factor was knocked down with siRNA, the cellular process altered drastically. The bCSCs morphology was changed, the cells were undergoing apoptosis; it was also found the expression of Oct-4 was downregulated. Most interestingly, Oct-4 knockdown not only influenced the cellular sustenance but also affected tumour formation ability. It was previously reported the Vitamin C causes apoptosis in bCSCs, and here we have shown that Vitamin C also interacts with Oct 4 by flow cytometry and sillico docking. In summary, Oct-4 can be a potential target for the CSC-based therapeutic maneuvering for breast cancer treatment.

To further understand the possible interaction between the Oct-4 and Vitamin C, Auto dock vina was used to investigate the binding sites and type of interactions occurring between Vitamin C and Oct-4 protein. Vitamin C – Oct-4 complexes were analysed using the molecular (blind) docking approach. Out of nine different predicted conformations for Vitamin C–Oct-4, top three ligand conformations having minimum binding energies were considered. The top minimum energies were as mentioned **-5.1, -5.0** and **-4.9 Kcal** mol^-1^ respectively. All the structural or binding parameters are summarized in Table 1. These results predicted that Vitamin C binds with Oct-4 mainly in beta position of K9 to K11 sequences. The major type of interaction between Vitamin C and Oct-4 is hydrogen bond between hydroxyl groups of the ligand to the oxygen or nitrogen atoms of amino acid main chain of the protein. The number of hydrogen bonds found for all the three different positions were seen in PyMol as shown in Fig. -4C-G.

## Discussion

Octamer-binding transcription factor 4 (Oct-4), a homeodomain transcription factor of the Pit-Oct-Unc (POU) family, is considered to play an essential role in the self-renewal epithelial-mesenchymal transition (EMT) (Zeineddine et al., 2014) and drug resistance development of CSCs and breast cancer metastasis (Wang et al., 2014). Oct-4 is abundantly proved as one of the most significant transcription factors that control self-renewal and pluripotency of pluripotent stem cells and malignant tumour succession and differentiation in CSCs (Wu & Schöler 2014). Hence, Oct-4 could be one of the critical factors that target stem cell-specific cancer therapeutics.

Breast cancer, which is now a frequently detected type of cancer, is still one of the most lethal malignancies despite recent advances in early detection and therapy (Bose et al., 2018). Conventional therapies such as immune, chemo, and radiotherapy, can target breast cancer cells. However, conventional therapies are often unaffordable and toxic to the patients (Keegan et al., 2012; Arruebo et al., 2011). Hence, certain alternatives/additives to conventional therapies such as oral supplements, nutraceuticals, and antioxidants might provide a better quality of life and the mitigation of the disease burden in breast cancer patients (Lopes, Dourado & Oliveira el., 2017). CSCs play critical roles in regulating tumour initiation, relapse, and chemoresistance, and hence such processes must be disrupted as therapeutic strategies (Lopes, Dourado & Oliveira el., 2017). In the current study, we have used the MDA MB 468 cell line as an in vitro TNBC model and isolated the CSCs based upon the surface markers CD44^+^/24^-^. The CD44^+^/24^-^ CSCs were further characterized for the validation of the CSC phenotype. We obtained a high expression of Oct-4 in the cancer CSC population by IF and flow-cytometric analysis (94%). Moreover, detection of Oct-4 in metastatic cancer cells and tissues indicates its enrichment CSCs (Kim, & Nam 2011).. Considerable uncertainties and controversies persist, and only a few studies have been reported attempting to target Oct-4 directly (Wang et al., 2015). According to the available reports, inhibition of Oct-4 effectively suppressed the propagation of human embryonal carcinoma cells and triggered their apoptotic death (Kim, & Nam 2011). This current study additionally talks about the dependence of bCSC on Oct-4 for their survival/ sustenance.

Nevertheless, our results indicate a negative co-relation between Oct 4 expression and apoptosis. Oct-4 is also known as an anti-apoptotic protein; hence repression of Oct-4 can cause apoptosis in breast cancer (Phi et al., 2018). This current study added to the growing evidence on the previous study’s conclusion wherein we demonstrated the cellular apoptosis of bCSCs upon Oct-4 knock-down. Furthermore, in this work, we have demonstrated the morphological changes and disruption of mammosphere formation of bCSCs upon si-RNA mediated Oct-4 knock-down in vitro. Thus, depending on the cellular contexts, restraining Oct-4 by siRNA may activate or inactivate its downstream counterparts, resulting in significant apoptosis in bCSCs, resulting in therapeutic outcomes. (Kristensen et al., 2010). Previous reports attributed the inhibition of Oct-4 to effectively suppress the propagation of human embryonal carcinoma cells and triggered their apoptotic death (Phi et al., 2018). This study established that Vitamin C in bCSC downregulates the Oct-4 expression, further leading to apoptosis as it is known that the Vitamin C increases cellular ROS levels. The interaction between Vitamin C and Oct-4 was finally confirmed by in Silico docking, which added novel information in the literature.

Moreover, we anticipate that Oct-4 knockout by CRISPR-Cas9 technology may attain a higher degree of inhibition on cell propagation than si-Oct-4 in bCSCs (Phi et al., 2018). Targeting major markers are known in the literature, where CD44 was knocked down in the bCSCs and reported the reduced stem ness of the bCSC, as a promising breast cancer therapy approach (Pham et al., 2011). Although, the abundant markers such as CD44, depletion of CD44 in other tissue can cause secondary damage, which can finally lead to failure in cancer therapy. Hence, the cancer specific therapy will be more promising. Such a strategy may help tackle the significant challenges in treating human cancers, which is worth exploring in next step. When specificity and comes, strategies like micro-RNA (miRNA) is one of the crucial components for the breast cancer prognosis (Ko et al.,2020). We established an essential proof-of-concept that inhibiting cancer-specific Oct-4 can effectively target CSCs and the entire bulk of cancer cells (Li et al., 2017). We, hence, propose down-regulation of Oct-4 first at a pre-clinical level in tumour xenograft animal models followed by possible clinical trials using either RNAi or CRISPR-Cas9 approaches.

## Conclusion

This study depicts the role of Oct-4 in the sustenance of the bCSCs. Oct-4 is, hence, one of the essential cancer-specific transcription factors; which gets downregulated upon treatment with Vitamin C leading to cellular apoptosis. in bCSCs. Further, we established the interaction between the Oct-4 and Vitamin C by the in-silico chemical docking technique. In addition, Oct-4 could be the best candidate target for future breast cancer therapeutics for its unique abundance in metastatic cancer and its contribution to its sustenance. Finally, the molecule Oct-4 can be a promising therapeutic target to improvise the lives of numerous breast cancer patients.

## Funding and Acknowledgements

The authors would like to thank Indian Council of Medical Research (ICMR) for providing the senior research fellowship to Mr Utsav Sen (Ph.D. scholar) [ICMR-2017-3769/CMB/BMS]. HKR thanks Council of Scientific & Industrial Research, Govt. of India for his doctoral fellowship. Authors also thank the Yenepoya Research Centre, Yenepoya (Deemed to be University) Mangalore for its infrastructural and administrative support for conducting this research. This is IICT communication number IICT/Pubs./2021/125

## Author contribution

US designed and performed the experiments, analysed, interpreted data, and wrote the initial manuscript, and prepared the figures; SS and MN carried out the experiment required in the revision of the manuscript, AG carried out in silico docking and its data interpretation; HKR developed the lipid transfection reagent and helped in siRNA experiment. RB provided his valuable inputs in the entire manuscript, provided cationic lipid delivery system, reviewed and approved the manuscript; SS and BB directed the project, involved in designing experiments and analysis of data, wrote and approved the manuscript.

## 7. Supporting information

Additional tables and information can be found in the Supporting information section.

